# Distraction attenuates goal-directed neural responses for food rewards

**DOI:** 10.1101/2020.01.13.904532

**Authors:** Iris Duif, Joost Wegman, Kees de Graaf, Paul A.M. Smeets, Esther Aarts

## Abstract

Distracted eating can lead to increased food intake, but it is unclear how. We hypothesized that distraction affects the change in motivated responses for food reward after satiation. To investigate this, 38 healthy normal-weight participants (28F, 10M) performed a detection task varying in attentional load (high or low distraction) during fMRI. Simultaneously, they exerted effort for food rewards (sweet or savory) by repeated button presses. Two fMRI runs were separated by outcome devaluation (satiation) of one of the reward outcomes, to assess outcome-sensitive, i.e. goal-directed, responses. Behavioral results showed no effect of distraction on effort for food reward following outcome devaluation. At an uncorrected threshold (*p*<0.001), distraction decreased goal-directed responses (devalued versus valued) in the right inferior frontal gyrus (rIFG). Importantly, these distraction-sensitive rIFG responses correlated negatively (*r* = - 0.40, *p* = 0.014) with the effect of distraction on the number of button presses. Specifically, decreased rIFG responses due to distraction related to increased button presses for food reward after satiation, in line with the rIFG’s established role in response inhibition. Furthermore, distraction decreased functional connectivity between the rIFG (seed) and left putamen for valued versus devalued food rewards (*p*FWE*(cluster)*<0.05). Our results suggest that distraction attenuates the ability to inhibit responses for food reward after satiation by affecting the rIFG. Furthermore, distraction attenuated connectivity between two regions involved in response inhibition – rIFG and putamen – after outcome devaluation. These results may explain why distraction can lead to overeating in our current, distracting, environment. The study was preregistered at: https://osf.io/ad2qk.

## Introduction

Multi-tasking with electronic devices, such as our smart phones or computers, has become common behavior in everyday life (Carrier et al., 2015), and increasingly occurs during consumption of food (de Graaf and Kok, 2010; Teo et al., 2018). Such “distracted eating” has been shown to cause overeating (Robinson et al., 2013), is associated with increased BMI (Bickham et al., 2013), and with increased choices of palatable foods (Shiv and Nowlis, 2004).

However, it is unclear how distraction increases food intake. Previously (<u>Duif et al., submitted</u>), we showed that distraction attenuates functional connectivity during tasting between a primary taste region in the insula and a secondary taste region in the OFC, and that lower sweetness activation of the insula under high distraction predicted increased subsequent food intake. However, this design did not allow for investigation of other processes than taste, such as food-related motivational or decision-making processes that may also be affected by distraction.

For example, in the act of grasping another crisp out of a bowl, attention to the TV, instead of to the crisp, may bias decision-making processes to the advantage of automatic and reflexive, i.e. habitual, choices. During learning, both humans and rodents form response-outcome (R-O) and stimulus-response (S-R) associations. When a response is made based on the outcome (reward) value of an action, the response is governed by R-O associations, and leads to goal-directed actions. On the other hand, habitual actions can be controlled by S-R associations, which have no associative link with the outcome value of the action. Habitual actions are relatively automatic and therefore less costly compared with goal-directed actions (see Balleine & O’Doherty, 2010 for a review). When consuming a food to satiation, the outcome value of the eaten food decreases (Rolls et al., 1983; de Wit et al., 2003; Fletcher et al., 2010), causing people to discontinue eating if they were to act in a goal-directed manner. However, we hypothesize that when distracted, goal-directed control is attenuated, causing people to continue grasping for another crisp in a habitual manner and ignoring the decreased outcome value of the food.

One way to investigate goal-directed versus habitual behavior, is through outcome devaluation. For example, Valentin, Dickinson, and O’Doherty, (2007) used tomato juice (savory) and chocolate milk (sweet) as stimuli and devalued participants on one of these drinks through satiation (i.e. sensory-specific satiation). Before and after the devaluation, participants had to choose one of two rewards in an instrumental choice paradigm. Using fMRI during this task, they found that the ventromedial prefrontal cortex (vmPFC) was involved in goal-directed responses for the juices. Other fMRI studies in humans also found the vmPFC to be involved in encoding reward predictions based on goal-directed R-O associations (Tanaka et al., 2008; de Wit et al., 2009; Gläscher et al., 2009), in line with its role in value-based decision-making (Rangel, 2013).

To test the effect of distraction on goal-directed responses for food reward, we adapted a similar outcome devaluation paradigm as used previously (Valentin et al., 2007; Hogarth et al., 2012; Morris et al., 2015). However, instead of a binary choice, participants had to exert effort to obtain a sweet or savory food reward. During this extended period of effort, participants were less or more distracted by a categorical visual detection task of varying attentional load (high, low distraction). After outcome devaluation of the sweet or savory reward, participants performed the same task again. Under high (versus low) distraction, we expected participants to exert less goal-directed effort, and to find less activation in the vmPFC. Due to the nature of the task (effort for reward outcomes instead of binary choice), we also tested the effect of distraction on devalued versus valued food rewards in other fronto-striatal regions, related to other motivational or cognitive control processes than value-based decision-making (secondary outcomes, see <u>preregistration</u>).

## Methods

### Participants

Thirty-nine right-handed healthy participants took part in this study and were recruited from a previous study of which the results are presented elsewhere (<u>Duif et al., submitted</u>). All participants were recruited on a voluntary basis. They gave written informed consent and were reimbursed for participation according to institutional guidelines of the local ethics committee (CMO region Arnhem-Nijmegen, the Netherlands, protocol nr. 2015-1928). Participants received financial compensation for participating in the study.

As a result of one drop-out (i.e. not completing the test session), the final sample size of the study was 38 (age range 19 – 37; mean age (± SD): 23.8 (3.6); 28 females; mean Body Mass Index (BMI, ± SD): 22.3 (2.01)).

### Screening

On a separate session prior to the test sessions of the current and previous study (<u>Duif et al., submitted</u>), subjects were screened for exclusion criteria. To be eligible to participate in the study, participants had to have a BMI within a range of 18.5 – 30.0, had to be within 18 – 35 years old at the time of the intake session (mean (SD) difference in weeks between intake session and current test session: 66.81 (10.26)), and right-handed. Exclusion criteria were current pregnancy; MRI-incompatibility; diabetes mellitus; history of hepatic, cardiac, respiratory, renal, cerebrovascular, endocrine, metabolic or pulmonary diseases; uncontrolled hypertension; eating, neurological, or psychiatric disorders; current strict dieting; restrained eating score ≥ 20 percent highest percentile (≥ 3.60 for males and ≥ 4.00 for females) on the Dutch Eating Behavior Questionnaire (DEBQ; Van Strien et al., 1986; Table 1); current psychological or dietary treatment; taste or smell impairments; use of neuroleptica or other psychotropic medication; food allergies relevant to the study, deafness, blindness, and sensori-motor handicaps; drug, alcohol or nicotine addiction; inadequate command of both Dutch and English, and a change in body weight of more than 5 kg in the past two months.

**Table 1.** Neuropsychological measurements

|  | Mean | Standard deviation | Minimum | Maximum |
| --- | --- | --- | --- | --- |
| BIS | 15.8 | 3.1 | 10 | 24 |
| BAS | 24.6 | 5.1 | 18 | 35 |
| BIS-11 | 69.1 | 5.3 | 59 | 80 |
| Kirby | 0.0070 | 0.0080 | 0.0002 | 0.0255 |
| BES | 24 | 4.8 | 16 | 33 |
| SFFQ-DHD | 54 | 13.0 | 24 | 75 |
| DEBQ |  |  |  |  |
| *Restraint* | 2.2 | 0.7 | 1.0 | 3.9 |
| *Emotional* | 2.2 | 0.6 | 1.2 | 3.4 |
| *External* | 3.3 | 0.5 | 2.6 | 4.3 |
| PFS | 35.3 | 9.3 |  |  |
| TFEQ |  |  |  |  |
| *Diet* | 7.2 | 3.7 | 1.0 | 18.0 |
| *Disinhibition* | 5.2 | 2.7 | 2.0 | 14.0 |
| *Hunger* | 6.1 | 2.9 | 0.0 | 12.0 |
| FFMQ | 81.9 | 9.0 | 60.0 | 101.0 |

*BIS/BAS*: Behavioural Inhibition System/Behavioral Approach System questionnaire; *BIS-11*: Baratt Impulsiveness Scale-11; *Kirby*: delayed reward discounting questionnaire; *BES*: Binge Eating Scale; *SFFQ-DHD*: Short Food Frequency Questionnaire, Dutch Healthy Diet; *DEBQ*: Dutch Eating Behaviour Questionnaire; *PFS*: Power of Food Scale; *TFEQ*: Three Factor Eating Questionnaire; *FFMQ*: Five Facet Mindfulness Questionnaire. Note, the DEBQ, BIS-11, BES, PFS, BIS/BAS, and Kirby were filled out on a separate test session prior to the current study. N=38.

### Procedure

Participants came to the laboratory for one experimental test session. To standardize participants’ hunger, they were instructed to eat a standardized meal three hours before the session (yogurt drink, strawberry flavor (200 gram (g)., 850 kilojoules, 6.0 g. protein, 30.0 g. carbohydrates, 6.0 g. fat; Breaker, Melkunie, Nijkerk, the Netherlands)). Three hours before and after consumption of the yogurt drink, participants were instructed to abstain from eating solid foods and from drinking sugared or sweetened drinks (but not water), and to refrain from alcohol use (24 hours) and neuroleptic or psychotropic drug use (7 days).

After completing an informed consent form and a recent food intake questionnaire, participants practiced the first calibration to balance reward probabilities of the effort task across participants (see *Calibration phase – determination of the reward probability*). Next, they were asked to choose their preferred sweet and salty reward (sweet: wine gums, M&Ms, or Skittles; salty: cocktail nuts (crusted peanuts), Pringles (original), or salty crackers (TUC, paprika)). They rated how hungry, full, and thirsty they felt, and how much they wanted and liked (during tasting) their chosen sweet and salty reward on 100-mm digital visual analogue scales (VAS) ranging from 0 (“not at all”) to 10 (“very (much)”). Following the ratings, participants learned which color (blue, green) would later cue which reward (salty, sweet) through stimulus-outcome learning (see *Stimulus-Outcome learning*). Next, they practiced the task (see below). After this, anthropometric measurements (BMI (weight(kg)/(height (cm) ^2^)) and waist-hip ratio (waist(cm)/hip(cm)) were taken. Subsequently, participants performed the task in two fMRI runs (pre- and post-devaluation runs) in the same session, separated by the outcome devaluation phase, which was performed outside the MR scanner. Twice during the task (after block 1 and 2 of the pre- and post-devaluation runs) participants rated how hungry, full, and thirsty they were on VASs. Directly after each run, they received 1/5 of their winnings and consumed the snacks. A high-resolution anatomical scan (T1, see *(f)MRI data acquisition*) was performed prior to the pre-devaluation task, and a resting state scan followed the post-devaluation task. After the post-devaluation run, participants rated hunger, fullness, thirst, wanting and liking once more, and filled out questionnaires (The Five Facet Mindfulness Questionnaire (FFMQ; Baer, Smith, Hopkins, Krietemeyer, & Toney, 2006), The Three-Factor Eating Questionnaire (TFEQ; Stunkard & Messick, 1985), and the Short Food Frequency Questionnaire – Dutch Healthy Diet index (SFFQ-DHD; van Lee et al., 2016); Table 1).

### Distraction task: categorical visual detection task

Participants performed a categorical visual detection task during fMRI scanning (Fig. 1). Each trial (total duration: 14-18 s) started with a fixation cross (jittered duration of 1-2 s, uniformly distributed), followed by an instruction screen (1 s), indicating the category of pictures for which the participant should count the targets (furniture, tools, or toys), and the speed of the trial (‘>’ for a slow trial, ‘>>>’ for a fast trial. For example, if the instruction screen stated: “category: furniture, >>>”, this meant participants needed to count stimuli in the category furniture, and the pictures would be presented at high speed. In order to keep visual stimulation equal for both trial types, a visual mask always followed a picture. The visual masks were scrambled versions of the stimulus pictures, to keep luminance equal. For the low speed trials, both pictures and visual masks were presented for 750 ms. For the high speed trials, pictures were presented for 75 ms, and the visual mask for 675 ms. Consequently, there were twice as many pictures and visual masks in the high speed trials relative to the low speed trials (12 vs. 6), thus, a higher attentional load. At the end of each trial, participants had to indicate how many target stimuli (i.e. a picture belonging to the instructed category) they had seen by answering a 3-alternative forced choice (3AFC) question on screen (“How many targets did you see?”) with their right hand. One of the three answering options was the correct answer, the other two options varied one or two digits from the correct answer. For example, if the correct answer was “2”, the answering options would be: [0, 1, 2], [1, 2, 3], or [2, 3, 4]. The order of the options on screen was shuffled to ensure participants could not prepare their answer motorically. If they answered within the 1500ms time limit for responding to the question, the selected answer was presented (500 ms) in bold letters. If participants exceeded the time limit, the text “TOO LATE” was presented (500 ms) and the task continued. Participants made responses using an MRI-compatible button box. Participants received no feedback on whether they answered the target question correctly.

**Figure 1.**
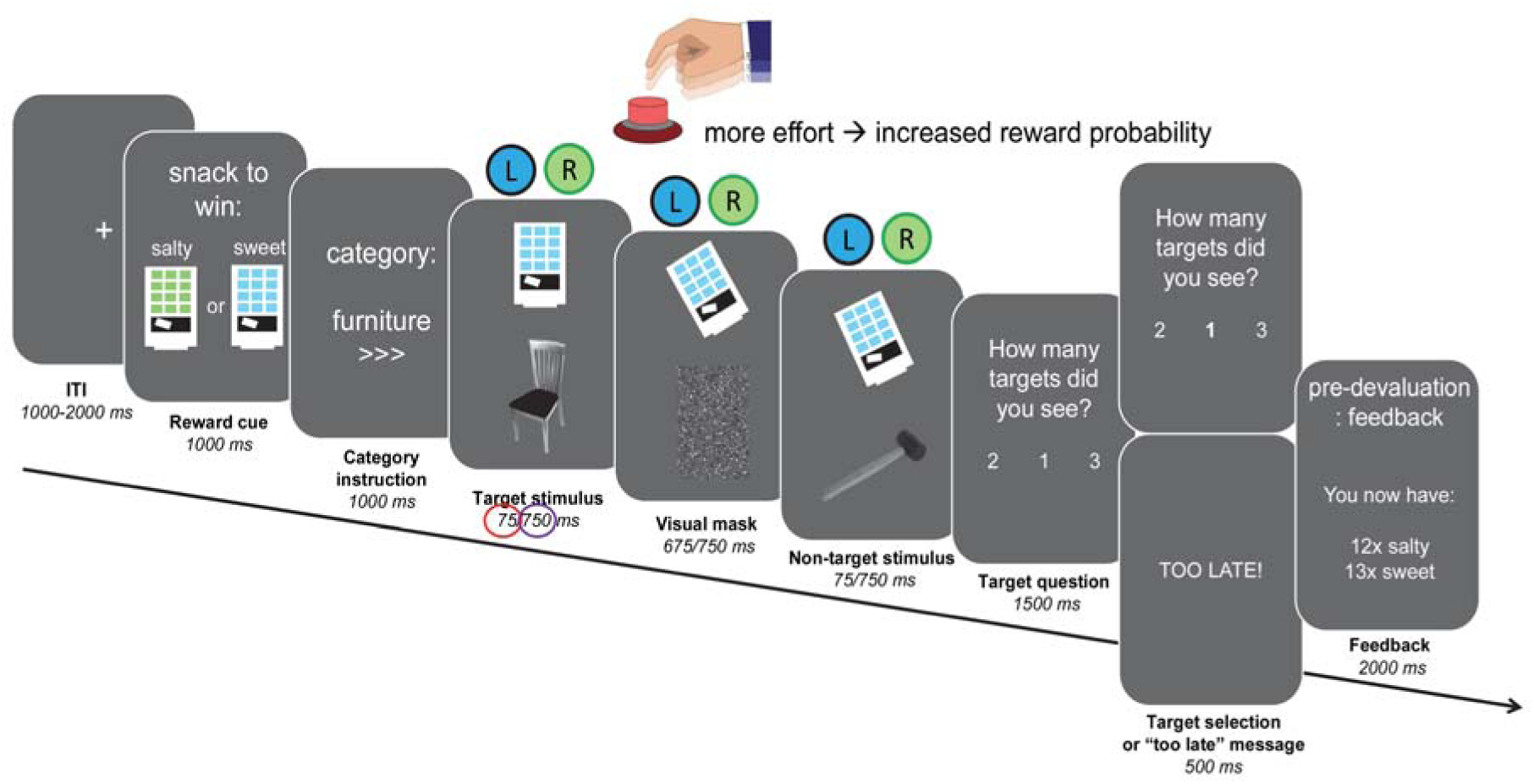
Trial structure for the categorical visual detection task with effort component. Each trial started with a verbal (word) and visual (color) cue indicating which snack could be won (word: sweet or salty, color: blue or green vending machine). An instruction screen followed, indicating the target category (furniture, tools, or toys) and difficulty (low (‘>’) or high (‘>>>’) of the trial. During the trial, target or non-target pictures were presented followed by a visual mask, and participants were instructed to count the pictures belonging to the instructed category. In the low distraction condition, six pictures were presented for 750 ms, giving subjects sufficient time to process the pictures. In the high distraction condition, twelve pictures were presented very briefly (only 75 ms), to increase the attentional load required to process the pictures. Simultaneously, a vending machine was presented on screen, which subjects could tilt by pressing one of two buttons with their left index finger to obtain the cued reward. Each of the two buttons was associated with either the sweet or the salty reward, which implied the vending machine could only be tilted during a trial when the button associated with the cued snack was used. The reward probability increased with the number of button presses subjects exerted (subjects have to exert effort to obtain the rewards), however never fell below 10% or above 90%. At the end of the trial, subjects indicated within 1500 ms how many target pictures they saw with their right hand (3AFC). Pre-devaluation, subjects received feedback on the amount of snacks won every 8 or 9 trials. The post-devaluation phase was performed in nominal extinction (i.e., no feedback was delivered).

### Effort task

During the distraction task, participants simultaneously performed the effort task to win sweet or salty rewards (Fig. 1). This task started with an instruction screen (1 s), which followed the instruction screen for the distraction task, and indicated which reward (salty, sweet) the participant could win during the trial. On screen, a vending machine (Morris et al., 2015) was presented with green or blue windows and the text: “snack to win: sweet”, or: “snack to win: salty”. The color of the vending machine was associated with the reward that could be won: the green color was always associated with winning the salty reward, and the blue color with the sweet reward (color-reward associations were counter-balanced). During the trial, the vending machine with the associated color was presented in the top middle of the computer screen, above the stimuli of the distraction task. To win the reward, participants had to exert effort to ‘tilt’ the vending machine to the left or right by repeatedly pressing the corresponding button (tilt left = left button, tilt right = right button) with their left index finger. The left button was always associated with obtaining one reward, the right with the other. Button-reward associations were counter-balanced across participants, and learnt by trial and error at task practice during which participants received direct feedback on whether they won the reward. During trials where the sweet reward could be won, the vending machine tilted only to the side associated with obtaining the sweet reward, and vice versa for the salty reward. Pressing the associated button more often resulted in a larger reward probability, i.e. exerting more effort related to a larger chance of winning the reward. Pressing the corresponding button at least once resulted in a reward probability of 10%. If participants pressed equally often or more often than the maximum number of button presses during the individual calibration (see: *Calibration phase – determination of the reward probability*), the reward probability on a trial would be 90%. Not pressing at all, or pressing any other button, resulted in a reward probability of 0%.

Post-devaluation, participants performed the dual task again during fMRI scanning. The pre- and post-devaluation tasks were similar, except no direct feedback was delivered on whether they won the reward post-devaluation (i.e. nominal extinction). Pre-devaluation, subjects received feedback on the amount of snacks won every 8 or 9 trials. Hence, these trials with feedback were longer (16-18 s instead of 14-15 s).

### Outcome devaluation phase

After participants performed the dual task for the first time in the MR-scanner (pre-devaluation run), they were taken to a behavioral lab for the outcome devaluation phase. Two bowls were placed in front of them. One bowl was filled with a fixed amount (10% of participants’ energy need, based on their basal metabolic rate (BMR, Schofield, Schofield & James, 1985)) of one of the snacks (counter-balanced), the other with an excessive amount of the same snack. Participants were asked to eat to satiation and to finish the fixed portion. If this portion did not satiate them, they were instructed to continue eating *ad libitum* from the other bowl. Unbeknownst to participants, we weighed the bowls before and after consumption to calculate the amount of calories consumed. Before and after the outcome devaluation phase, participants again rated how much they wanted and liked (after tasting) both rewards.

### Dual task: effort + distraction

Pre- and post-devaluation, participants performed in total eight blocks of 25 trials (a total of 200 trials), i.e. four blocks per run. Both pre- and post-devaluation runs had the following numbers of trials: 50 low distraction trials (25 salty reward, 25 sweet reward); 50 high distraction trials (25 salty reward, 25 sweet reward). During task practice, participants performed a total of 32 trials (6 trials: distraction task only; 6 trials: effort task only, 20 trials: full dual task (distraction + effort task)). Category and reward presentation were pseudo-randomized, i.e. the same category and reward were never presented more than 3 times in a row. Moreover, maximally two target stimuli were presented after another.

### Calibration phase – determination of the reward probability

Pilot data showed large inter-individual baseline differences in the number of button presses individuals can exert in a certain time frame. Therefore, prior to the pre-devaluation run, we measured participants’ individual maximum button presses in a calibration phase. Whilst in the MR-scanner, participants pressed each of the buttons (left and right) on the button box (used during the effort task) with their left index finger as often as they could during 14 seconds (the approximate duration of a trial in the dual task). The probability of obtaining a reward during a trial in the dual task was calculated based on the number of button presses participants exerted in the current trial, and their maximum number of button presses. The following formula was used in the calculation:

**Reward probability of a trial = minimum probability + ((maximum probability – minimum probability) * (current number of button presses in a trial / maximum number of button presses determined during calibration phase)).**

The minimum probability was set to 0.1, given that participants pressed the button corresponding to the reward at least once. The maximum probability was set to 0.9.

### Stimulus-Outcome learning

To ensure participants learned the associations between the color (stimuli: blue, green) of the vending machine and the rewards (outcomes: sweet, salty), we provided them with three pieces of their preferred sweet and salty reward and paired each piece with digital presentations of the associated color. On screen, a blue or green square was presented (3 s) which was followed by the text: “Now, please eat one of the [X] snacks. Press ENTER on the keyboard after you’ve finished eating it”. [X] was filled with “SWEET” or “SALTY”. In total, 6 stimulus-outcome pairings were presented (3 sweet, 3 salty). Prior to each trial in the actual fMRI task, i.e. at the time of the instruction screen cueing the reward that could be won during the trial, the trial-specific color-reward association was repeated. At the end of the test session, we asked participants to recall the associations.

### Behavioral analyses: performance

To test whether participants’ performed significantly better on the low (versus high) load trials of the distraction task (i.e. correct detection of the number of targets in a trial), we calculated mean weighed accuracies across trials per condition using the following formula: **weighed accuracy = 1 – (mean(abs(actual number of targets – chosen number of targets))).** In this calculation, the absolute difference (abs, i.e. the non-negative value of the difference without regard to its sign) between the correct answer and the answer participants gave was used to account for whether participants were 1 or 2 targets off the correct answer (the latter case being more erroneous than the former). Mean weighed accuracies were analyzed using repeated measures ANOVA (IBM SPSS Statistics 23, Chicago, IL) with attentional Load (high, low), Reward (valued, devalued), and Time (pre-, post-devaluation) as within-subject factors.

### Behavioral analyses: devaluation effect

As a subjective measure of devaluation magnitude, we assessed wanting ratings for the sweet and salty snack over time (measured at four time points: at baseline (t_0_), directly before the devaluation (t_1_), directly after the devaluation (t_2_), and after completing the post-devaluation run (t_3_)). If the devaluation was successful, we expected the wanting ratings to decrease significantly for the reward that would be devalued, and not for the reward that would remain valued. This was tested using repeated measures ANOVA with within-subject factors Time (t_0_, t_1_, t_2_, t_3_) and Reward (valued, devalued). Additionally, we tested for pre-experimental wanting differences (wanting at t_0_) for the valued versus devalued snack using a paired-samples *t*-test.

### (f)MRI Data Acquisition

To measure blood oxygen level dependent (BOLD) contrast, whole-brain functional MRI images were acquired on a Siemens 3T Skyra MRI scanner (Siemens Medical system, Erlangen, Germany) using a 32-channel head coil. During the task, 3D echo planar imaging (EPI) scans using a T_2_*weighted gradient echo multi-echo sequence (<u>Centre for Magnetic Resonance Imaging, University of Minnesota</u>) were acquired (voxel size 3.5 x 3.5 x 3 mm, TR = 2070 ms, TE = 9 ms; 19.1 ms; 29.2 ms; 39.3 ms, FoV = 224 mm). The slab positioning and rotation (approximate average angle of 14 degrees to AC axis) optimally covered both prefrontal and subcortical brain regions. Before the acquisition of functional images, a high-resolution anatomical scan was acquired (T_1_-weighted scan, MPRAGE, voxel size 1×1×1 mm, TR = 2300 ms, TE = 3.03 ms, 192 sagittal slices, flip angle 8°, field of view 256 mm).

### (f)MRI Image Processing

Data were analyzed using SPM8 (www.fil.ion.ucl.ac.uk/spm) and FSL version 5.0.11 (http://www.fmrib.ox.ac.uk/fsl/). The volumes for each echo time were realigned to correct for motion artefacts (estimation of the realignment parameters is done for the first echo and then copied to the other echoes). The four echo images were combined into a single MR volume using an optimized echo weighting method (TE-weighting (Poser et al., 2006; Posse, 2012)). Combined functional images were slice-time corrected by realigning the time-series for each voxel temporally to acquisition of the middle slice and spatially smoothed using an isotropic 8 mm full-width at half-maximum Gaussian kernel. Next, ICA-AROMA (non-aggressive; Pruim et al., 2015) was used to reduce motion-induced signal variations in the fMRI data. Subject-specific structural and functional data were then coregistered to a standard structural or functional stereotactic space (Montreal Neurological Institute (MNI) template) respectively. After segmentation of the structural images using a unified segmentation approach, structural images were spatially coregistered to the mean of the functional images. The resulting transformation matrix of the segmentation was then used to normalize the anatomical and functional images into MNI space. The functional images were resampled to 2 x 2 x 2 mm using trilinear interpolation.

### Statistical fMRI analysis

Statistical analysis of fMRI data was performed using a general linear model (GLM) approach. The images of both experimental runs (pre- and post-devaluation runs) were combined into one model. At the individual (first) level, subject-level data were analyzed using a fixed effects model, which included four regressors of interest per run. These modeled the trials of valued reward, low load; valued reward, high load; devalued reward, low load; devalued reward, high load. Durations of these regressors represented the moment the first picture of the detection task was presented until presentation of the target question (mean duration: 9.18 s, SD: 0.09 s, SEM: 0.001 s). Parametric modulators reflecting the number of button presses per trial were added to each regressor of interest, to correct for signal change induced by the difference in number of button presses between the low and high load condition. Six additional regressors of non-interest were added, representing onsets of the reward instruction screen (one for valued and one for devalued reward trials), and presentation of the target question (“How many targets did you see?”, one for low and one for high load trials). The other two regressors reflected average signal variation in white matter and cerebrospinal fluid regions. All regressors were convolved with the canonical hemodynamic response function. High pass temporal filtering (128 s) was applied to the time series of the functional images to remove low-frequency drifts and correction for serial correlations was done using an autoregressive AR(1) model.

At the group level, we assessed the main effect of Load (high > low load) by contrasting high with low load trials across both runs (pre, post) and rewards (valued, devalued). To identify brain areas sensitive to goal-directed effort, we computed the difference between the two rewards post-devaluation (valued, devalued) relative to their respective baseline (pre-devaluation) during button presses for reward.

To identify our *a priori* defined vmPFC region in this functional contrast (see <u>preregistration</u>), we used an 8-mm spherical region-of-interest (ROI) centered on coordinates [-2, 32, −21] derived from averaged coordinates from previous studies that reported vmPFC responses during instrumental conditioning and/or outcome devaluation (Daw et al., 2006; Kim et al., 2006; Valentin et al., 2007). We used the Hammersmith atlas to determine whether the activated areas overlapped with the fronto-striatal regions defined in the atlas (bilateral caudate nucleus (regions 34;35), nucleus accumbens (36;37) and putamen (38;39) for striatum, bilateral middle frontal gyrus (28;29), Hammers et al., 2003), precentral gyrus (50:51), straight gyrus (52;53), anterior orbital gyrus (54;55), inferior frontal gyrus (56;57), superior frontal gyrus (58;59), medial orbital gyrus (68;69), lateral orbital gyrus (70;71), posterior orbital gyrus 72;73), subgenual frontal cortex (76;77), subcallosal area (78;79), and pre-subgenual frontal cortex for the frontal regions (Gousias et al., 2008).

To investigate the effect of Load on processing in the vmPFC and other fronto-striatal regions involved in goal-directed control, we assessed the interaction effect of Load (low > high Load) x Reward (valued > devalued) x Time (post > pre). The activated regions for the goal-directed effort contrast (i.e., Time x Reward) were used as ROIs for the Load x Reward x Time interaction contrast. Mean beta weights were extracted from all voxels in the identified ROIs using MarsBar (Brett et al., 2002). These beta-weights were analyzed using ANOVA with the same factors as in the whole-brain analyses (Load, Reward and Time).

### Distraction-related functional connectivity analysis

We used a generalized psychophysiological interaction (gPPI; McLaren, Ries, Xu, & Johnson, 2012) analysis to investigate whether the (seed) region(s) showing a distraction effect on goal-directed control (Load x Reward x Time) exhibited differential fMRI BOLD connectivity for the same 3-way interaction. To estimate the neural activity producing the physiological effect in the seed region for each subject, the BOLD signal was extracted from this region and deconvolved (Gitelman et al., 2003). This was included in the model as the physiological regressor. The task regressors for each of the relevant task conditions (i.e. the psychological regressors: low load, valued reward; low load, devalued reward; high load, valued reward; and high load, devalued reward) were also added to the model. The psychophysiological interaction was entered by multiplying the estimated neural activity (i.e., the physiological regressor) by the duration times for each of the task conditions (i.e., the psychological regressors) separately convolved with the HRF, resulting in nine regressors of interest per run on the first level (i.e., one physiological, four psychological, and four interaction regressors). For each participant, we created a PPI contrast for the 3-way interaction effect of Load (high>low load), Reward (valued>devalued reward), and Time (post>pre devaluation). On the group level, this PPI contrast was analyzed separately using a one-sample *t*-test.

All fMRI analyses were performed on the whole-brain level and with use of ROI analysis where specified. The results of all random effects fMRI analyses were thresholded at *p* < 0.001 (uncorrected) and statistical inference was performed at the cluster level, family-wise-error-corrected (*p*FWE < 0.05) for multiple comparisons over the search volume (the whole brain or a smaller search volume based on the defined ROIs).

### Effect of distraction on goal-directed effort (repeated button presses)

On each trial of the effort task, participants tilted the vending machine a number of times through button presses, i.e. they exerted effort, to obtain the associated reward. Mean effort values were calculated by averaging the number of button presses across all trials for each condition. To test whether effort decreased significantly for the devalued, but not for the valued reward (i.e. the degree to which effort became goal-directed after devaluation), we assessed the Time (pre, post) x Reward (valued, devalued) interaction using repeated measures ANOVA. The effect of distraction on goal-directed effort was determined by looking at the Load (high, low) x Reward (valued, devalued) x Time (pre, post) interaction.

### Analyses: brain-behavior correlations

Exploratory, we investigated whether the effect of Load on goal-directed effort covaried significantly with the effect of Load on processing in areas sensitive to goal-directed control. We executed a repeated measures ANCOVA with Load (high, low), Reward (valued, devalued), and Time (pre, post) as within-subject factors. The (brain) effect of Load in areas sensitive to goal-directed control reflected in the averaged extracted beta weights was used as dependent variable, and the difference score for the same effect on (behavioral) effort as covariate.

### Additional analyses: liking ratings

Similar to the analysis of the wanting ratings, we tested whether mean liking ratings for the valued and devalued reward changed significantly over time (using RM-ANOVA with Reward (valued, devalued) and Time (t_0_, t_1_, t_2_, t_3_) as within-subject factors). In addition, we tested for baseline differences (t_0_) in liking of the valued versus devalued reward using a paired-samples t-test.

### Additional analyses: hunger, fullness and thirst ratings

Hunger, fullness, and thirst ratings were completed 8 times during the experiment (at baseline (t0), after block 1 and 2 of the task (pre: t1, t2; post: t5, t6), directly before the devaluation (t3), directly after the devaluation (t4), and after completing the post-devaluation run (t7)). Using ANOVA, we tested whether hunger decreased and fullness increased after the devaluation. Participants were allowed to drink water during the outcome devaluation phase to ensure participants did not stop eating due to feelings of thirst. Therefore, we did not have specific hypotheses for changes in thirst.

### Additional analyses: hierarchical Regression

As a secondary analysis, we performed hierarchical regression analyses (stepwise method using backward elimination) to determine effects of individual differences. The effects of distraction on goal-directed effort (button presses) and those on processing of fronto-striatal areas involved in goal-directed control were used as dependent variables. Wanting for the two rewards (valued and devalued, directly before versus directly after the devaluation (t_4_-t_3_)) was added as independent variable to the first level. To the second level, liking of the two rewards (valued and devalued, directly before versus directly after the devaluation (t_4_-t_3_)), performance on the working memory task, hunger, fullness, and thirst ratings (t_4_-t_3_), and the following questionnaires were added: DEBQ, Baratt Impulsiveness Scale (BIS-11; Patton, Stanford, & Barratt, 1995), Binge Eating Scale (BES; Gormally, Black, Daston, & Rardin, 1982), Power of Food Scale (PFS; Lowe et al., 2009), Behavioral Inhibition/Activation System (BIS/BAS; Carver & White, 1994), Kirby monetary choice questionnaire (Kirby, 2009), FFMQ, TFEQ, SFFQ-DHD), see Table 1. Note, the DEBQ, BIS-11, BES, PFS, BIS/BAS, and Kirby were filled out on a separate test session prior to the current study.

## Results

### Behavioral results: performance on the distraction task

To test whether our load manipulation to manipulate distraction (Fig. 1) worked, we compared performance between the high and low load conditions. Indeed, participants were less accurate when answering the 3AFC target question (“How many targets did you see?”) when the visual targets were rapidly presented, i.e. the high load trials (weighed mean accuracy (± SEM): 0.63 (0.02)), than when the visual targets were slowly presented, i.e. the low load trials (weighed mean accuracy (± SEM): 0.81 (0.02)), (*F*(1,37) = 68.31, *p* < 0.001). Reward outcome or time did not affect performance (Reward (valued, devalued): *F*(1,37) = 1.23, *p* = 0.27; Time (pre, post): *F*(1,37) <1, *p* = 0.63, Table 2).

**Table 2.** Effort and weighed task accuracy

| | valued | | devalued | | $p$ | $F$ |
| --- | --- | --- | --- | --- | --- | --- |
|  | pre | post | pre | post |  |  |
| *Effort* |  |  |  |  |  |  |
| Low distraction | 39.4(0.9) | 39.9(1.0) | 39.2(0.9) | 39.4(1.1) | 0.61 | < 1 |
| High distraction | 40.4(0.9) | 40.7(1.0) | 40.0(0.9) | 40.1(1.3) | 0.57 | < 1 |
| *Weighed accuracy* |  |  |  |  |  |  |
| Low distraction | 0.80(0.03) | 0.82(0.02) | 0.79(0.02) | 0.82(0.02) | 0.65 | < 1 |
| High distraction | 0.61(0.03) | 0.62(0.03) | 0.65(0.03) | 0.63(0.02) | 0.48 | < 1 |

Means and standard errors are shown for effort (number of button presses exerted to obtain the rewards) and weighed task accuracy, pre- and post-devaluation, for each reward outcome (valued, devalued) separately, and Time (pre, post) x Reward (devalued, valued) statistics.

### Behavioral results: wanting ratings and effort for reward

We assessed the change in wanting ratings over time for the valued versus devalued reward, as a measure of the devaluation magnitude induced by satiation on the (to be) devalued reward. As expected, how much participants wanted the snack decreased significantly for the devalued, but not for the valued, reward after devaluation (Time (t_0_, t_1_, t_2_, t_3_) x Reward (valued, devalued): *F*(1,35) = 31.00, *p* < 0.001, Table 3). We further found main effects of Time (*F*(1,35) = 31.82, *p* < 0.001) and Reward (*F*(1,37) = 33.86, *p* < 0.001), reflecting an overall decrease in wanting for both rewards after the devaluation, and overall lower wanting ratings for the devalued relative to the valued reward. At baseline (t_0_), wanting did not differ significantly between the rewards (*t*(1,37) = 1.56, *p* = 0.13). Thus, the devaluation manipulation was successful. Similar results were found for changes in liking ratings (*see: Liking ratings*).

**Table 3.** Self-reported hunger, fullness, and thirst ratings, and wanting and liking ratings for each food reward (valued, devalued).

| | $t_{(0)}$ | $t_{(1)}$ | $t_{(2)}$ | $t_{(3)}$ | $t_{(4)}$ | $t_{(5)}$ | $t_{(6)}$ | $t_{(7)}$ | $p$ | $F$ |
| --- | --- | --- | --- | --- | --- | --- | --- | --- | --- | --- |
| *Hunger, fullness, and thirst ratings* |  |  |  |  |  |  |  |  |  |  |
| Hunger | 7.2(0.3) | 7.5(0.3) | 7.3(0.4) | 6.1(0.4) | 2.8(0.4) | 3.8(0.4) | 3.7(0.4) | 3.6(0.4) | <0.001 | 24.20 |
| Fullness | 1.4(0.2) | 1.5(0.2) | 1.5(0.2) | 3.0(0.3) | 7.1(0.3) | 5.9(0.4) | 5.9(0.4) | 5.6(0.4) | <0.001 | 68.98 |
| Thirst | 4.6(0.4) | 5.8(0.4) | 6.2(0.4) | 4.0(0.4) | 5.0(0.3) | 5.0(0.3) | 5.0(0.3) | 4.7(0.3) | <0.001 | 6.17 |
| *Wanting for each reward* |  |  |  |  |  |  |  |  |  |  |
| Valued | 6.7(0.4) | - | - | 6.4(0.4) | 5.9(0.4) | - | - | 6.3(0.4) | 0.40 | 1.01 |
| Devalued | 6.0(0.5) | - | - | 6.7(0.4) | 1.8(0.3) | - | - | 2.9(0.3) | <0.001 | 41.00 |
| *Liking for each reward* |  |  |  |  |  |  |  |  |  |  |
| Valued | 7.4(0.3) | - | - | 7.4(0.3) | 6.9(0.3) | - | - | 7.2(0.3) | 0.26 | 1.41 |
| Devalued | 8.2(0.2) | - | - | 7.6(0.3) | 4.4(0.4) | - | - | 5.1(0.3) | <0.001 | 38.29 |

Means and standard errors per time point, and Time statistics are shown.

Furthermore, we tested whether the devaluation effect – reflected in the Time (pre, post) x Reward (valued, devalued) interaction – appeared in the number of button presses exerted to obtain the rewards (Table 2). This interaction was not significant (*F*(1,37) < 1, *p* = 0.57). However, we did find a significant main effect of Load (*F*(1,37) = 38.59, *p* < 0.001). Participants pressed more often in the high versus low load condition (M_highload_ (± SEM): 40.31 (0.95), M_lowload_ (± SEM): 39.48 (0.95)). Furthermore, we found a trend main effect of Reward (*F*(1,37) = 3.00, *p* = 0.09). Participants tended to press more often for the valued than for the devalued reward (M_valued_ (± SEM): 40.11 (0.90), M_devalued_ (± SEM): 39.68 (0.96)). Finally, we tested whether attentional load affected goal-directed effort for the food rewards (i.e., as a function of outcome devaluation). The interaction between Load, Time, and Reward was not significant (*F*(1,37) < 1, *p* = 0.88).

### fMRI results: effect of attentional load

To test whether our distraction manipulation – operationalized by varying attentional load – activated brain regions typically involved in such tasks, we first assessed the effect of Load (high>low) on a whole-brain corrected threshold (*p*FWE < 0.05, at the cluster-level). At this threshold, we indeed found an effect of attentional load (high load > low load across reward types and time) in fronto-parietal and visual regions when contrasting the high with low load conditions (Table 4, Fig. 2).

**Figure 2.**
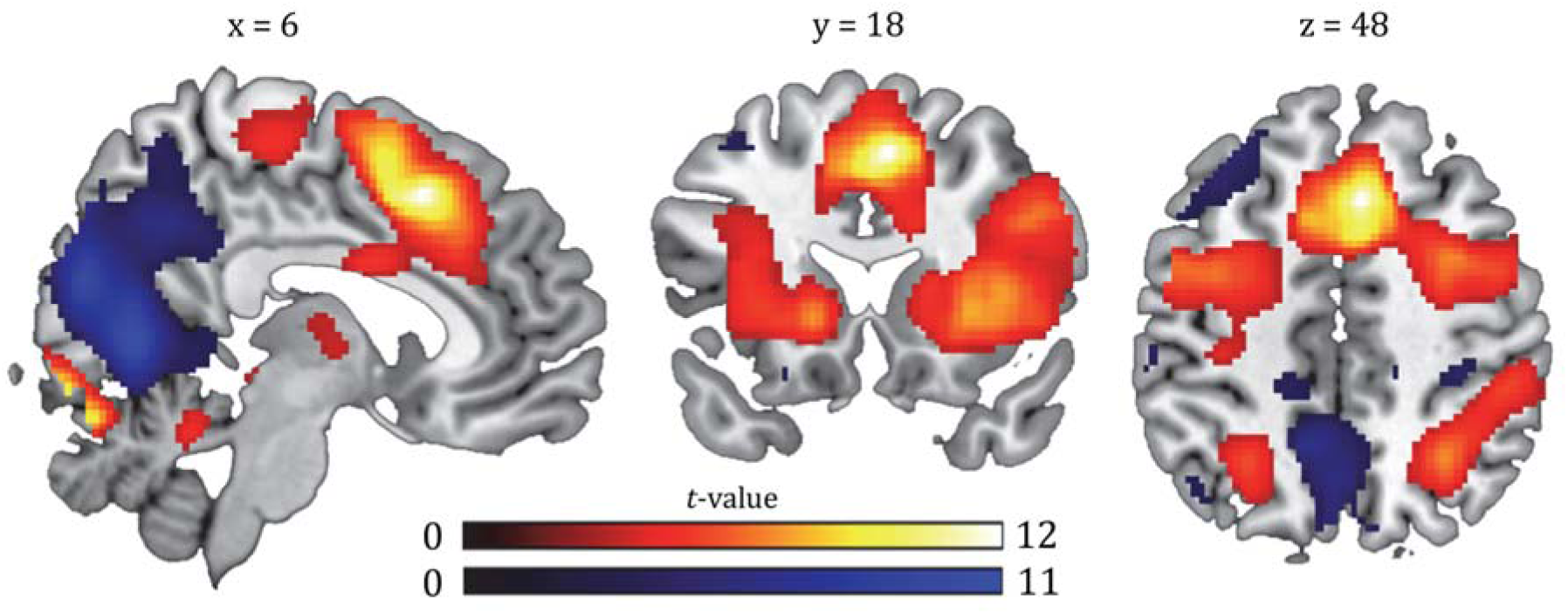
Main effect of attentional load (distraction manipulation). Contrast of high versus low (red, respectively), and low versus high load (blue, respectively) trials depicted at a *p* < 0.001, uncorrected threshold. For whole-brain (FWE < 0.05) corrected effects see Table 4. All statistical parametric maps were overlaid onto a T1-weighted canonical image (ch2better.nii.gz atlas using MRIcron software (http://www.mricro.com/mricron/install.html)). Slice coordinates were defined in MNI152 space and images are shown in neurological convention (left = left).

**Table 4.** Summary of brain regions exhibiting main effects of attentional load and time, and/or interactions with attentional load, reward, and time at a whole-brain corrected (FWE) cluster threshold. For uncorrected and small volume corrected effects see Fig. 2-5.

|  | Side (Left/Right) | MNI-coordinates |  |  | Size | *pFWE* | *t*-value |
| --- | --- | --- | --- | --- | --- | --- | --- |
| Label |  | x, y, z (mm) |  |  | (number of voxels) | (cluster-level) | (peak) |
| Main effect Attentional load, high > low load |  |  |  |  |  |  |  |
| Superior frontal gyrus | R | 6 | 18 | 48 | 17815 | < 0.001 | 12.55 |
| (Medial) Superior frontal gyrus |  | 0 | 8 | 58 |  |  | 11.22 |
| Inferior frontal gyrus | R | 34 | 28 | 10 |  |  | 9.90 |
| Lingual gyrus | R | 4 | -84 | -10 | 1315 | <0.001 | 9.96 |
| Cerebellum | R | 6 | -78 | -18 |  |  | 9.35 |
| Lingual gyrus | R | 12 | -92 | -2 |  |  | 8.56 |
| Superior parietal gyrus | R | 30 | -54 | 46 | 1320 | <0.001 | 7.28 |
| Supramarginal | R | 50 | -34 | 42 |  |  | 6.52 |
| Precentral gyrus | L | -10 | -22 | 66 | 1027 | <0.001 | 6.68 |
| Precentral gyrus | R | 4 | -24 | 62 |  |  | 5.21 |
| Precentral gyrus | R | 10 | -14 | 72 |  |  | 4.18 |
| Superior parietal gyrus | L | -26 | -54 | 44 | 788 | <0.001 | 6.30 |
| Medial occipital lobe | L | -28 | -68 | 32 |  |  | 4.73 |
| Inferior parietal lobe | L | -40 | -42 | 42 |  |  | 4.46 |
| Posterior temporal lobe | R | 66 | -34 | 12 | 392 | 0.003 | 5.58 |
| Posterior temporal lobe | R | 54 | -42 | 10 |  |  | 5.50 |
| Superior temporal gyrus, posterior part | R | 50 | -24 | -2 | 267 | 0.017 | 4.70 |
| Main effect Attentional load, low > high load |  |  |  |  |  |  |  |
| Lingual gyrus | R | 12 | -66 | 6 | 18096 | <0.001 | 11.08 |
| Cuneus | L | -4 | -66 | 14 |  |  | 10.97 |
| Cuneus | R | 12 | -76 | 20 |  |  | 10.38 |
| Precentral gyrus | L | -42 | 0 | 8 | 281 | 0.014 | 6.34 |
| Supramarginal | L | -64 | -30 | 26 | 550 | <0.001 | 6.13 |
| Postcentral gyrus | L | -56 | -24 | 44 |  |  | 5.75 |
| Posterior temporal lobe | L | -44 | -62 | 18 | 1197 | < 0.001 | 5.87 |
| Middle frontal gyrus | L | -28 | 32 | 50 | 216 | 0.039 | 5.60 |
| Insula | R | 38 | -10 | 4 | 221 | 0.036 | 5.56 |
| Rolandic operculum | R | 40 | -30 | 24 |  |  | 3.72 |
| Angular | L | 50 | -58 | 24 | 358 | 0.005 | 4.62 |
| Angular | R | 46 | -70 | 38 |  |  | 4.07 |
| Main effect of Time (post > pre) |  |  |  |  |  |  |  |
| Inferior parietal lobe | R | 34 | -52 | 44 | 1411 | <0.001 | 6.61 |
| Superior occipital lobe | R | 28 | -68 | 38 |  |  | 5.71 |
| Angular | R | 44 | -66 | 38 |  |  | 5.13 |
| Main effect of Time (pre>post) |  |  |  |  |  |  |  |
| Lingual gyrus | R | 18 | -70 | -12 | 325 | 0.004 | 5.15 |
| Cerebellum | R | 32 | -66 | -18 |  |  | 4.25 |
| Superior occipital lobe | R | 16 | -90 | 18 | 187 | 0.043 | 4.97 |
| Superior temporal lobe | L | -60 | -22 | 14 | 225 | 0.022 | 4.76 |
| Interaction effect of Reward x Load: devalued (low>high) > valued (low>high) |  |  |  |  |  |  |  |
| Supramarginal | R | 54 | -38 | 46 | 364 | 0.005 | 4.56 |
| Inferior parietal lobe | R | 44 | -38 | 30 |  |  | 4.44 |
| Inferior parietal lobe | R | 48 | -28 | 34 |  |  | 3.72 |
| Interaction effect of Time x Reward: post (valued>devalued) > pre (valued>devalued) |  |  |  |  |  |  |  |
| Superior frontal gyrus | R | 8 | 16 | 40 | 505 | 0.001 | 4.76 |
| gPPI – Interaction effect of Time x Reward x Load:<br>post (valued (low>high) > devalued (low>high) > pre (valued (low>high) > devalued (low>high))) |  |  |  |  |  |  |  |
| Thalamus | L | -22 | -16 | 0 | 485 | 0.001 | 5.39 |

### fMRI results: regions involved in goal-directed effort for food reward

To identify brain regions involved in goal-directed effort for the valued versus devalued reward, we assessed baseline-corrected differential responses for these rewards (Time(pre, post) x Reward(valued, devalued) interaction). The valued>devalued contrast showed significant whole-brain responses of a region in the anterior cingulate cortex (ACC: [8, 16, 40], *p*FWE(cluster) = 0.001, *t* = 4.76, k = 505, Fig. 3 (left), Table 4). In the devalued>valued contrast, on a *p* < 0.001 uncorrected threshold, we found increased responses in the vmPFC ([6, 52, −8], Fig. 3 (right)), which was our *a priori* defined search region. However, these were not significant after small volume correction using the spherical ROI described above (see *Methods*; *p*FWE(cluster) > 0.05).

**Figure 3.**
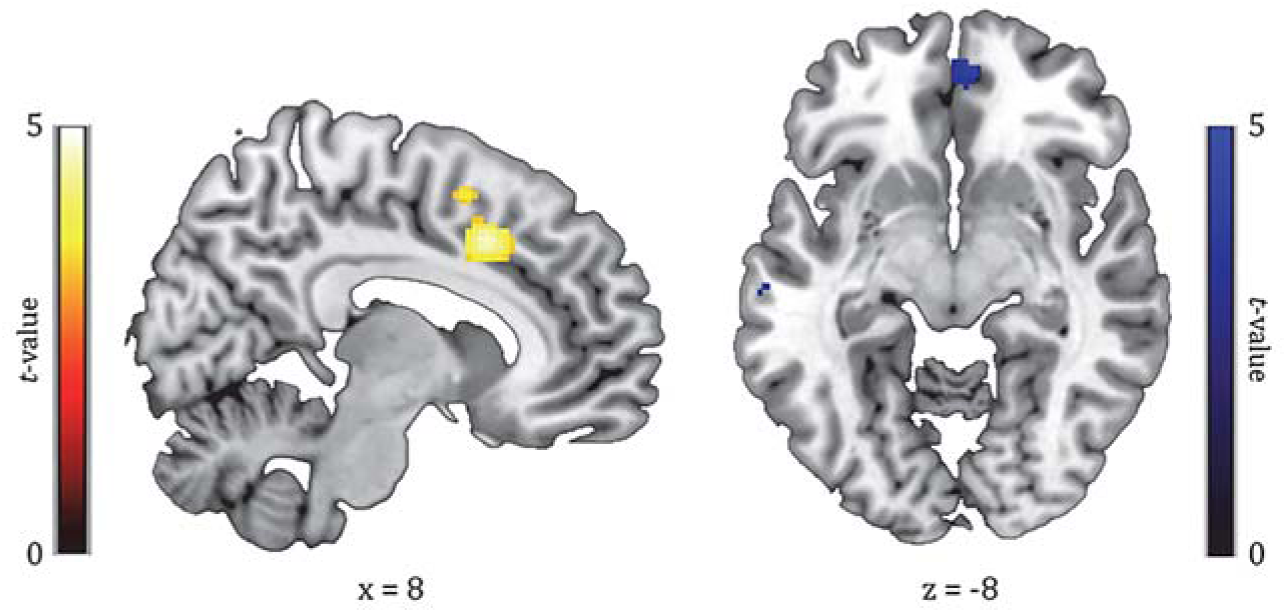
Time (pre,post) x Reward (valued, effect) interaction effect (manipulation of goal-directed effort), showing responses of the ACC (left; *p*FWE (cluster) = 0.001) and vmPFC (right; *p* < 0.001, uncorrected). Left (yellow): contrast of valued outcome versus devalued outcome trials (val>dev). Right (blue): contrast of devalued outcome versus valued outcome trials (dev>val). All parametric maps are depicted at a *p* < 0.001, uncorrected threshold.

### fMRI results: effect of attentional load on goal-directed effort for food reward

We assessed the effect of load on processing related to goal-directed control for food rewards in our ACC and vmPFC regions, using ROI analysis. The averaged extracted beta weights across the ACC cluster showing the effect related to goal-directed control (Time x Reward), did not show an interaction with Load (*F*(1,37) < 1, *p* = 0.52). The results did show a main effect of Load (*F*(1,37) = 87.98, *p* < 0.001), reflecting overall more ACC activity for the high, relative to low, load trials. The extracted beta weights across the vmPFC cluster for Time x Reward also showed no interaction with Load (*F*(1,37) < 1, *p* = 0.53), but did show a main effect of Reward (*F*(1,37) = 6.72, *p* = 0.01) independent of Time, reflecting generally larger vmPFC responses for the devalued than the valued reward.

Interestingly, on a whole-brain p < 0.001 uncorrected threshold, a region in the right inferior frontal gyrus (rIFG) responded ([44,36,-2], *k* = 88, Fig. 4 (Panel A-C)) to the three-way interaction contrast of Load (low>high), Time (post>pre), and Reward (devalued>valued). This cluster was not significant after small volume correction with our bilateral fronto-striatal mask (*p*FWE*(cluster)* = 0.19, *t* = 4.29). We exploratively investigated whole-brain simple effects and found this region in the Time(post>pre) x Reward(devalued>valued) interaction under low ([48, 36, 0], *k* = 122, *p*FWE*(cluster)* = 0.11, *t* = 4.31, Fig. 4 (Panel A)), but not high load. Thus, distraction tended to attenuate responses of the rIFG for devalued versus valued food reward after outcome devaluation. Given this effect, we used the rIFG cluster in further connectivity and brain-behavior analyses (see <u>preregistration</u> and below).

**Figure 4.**
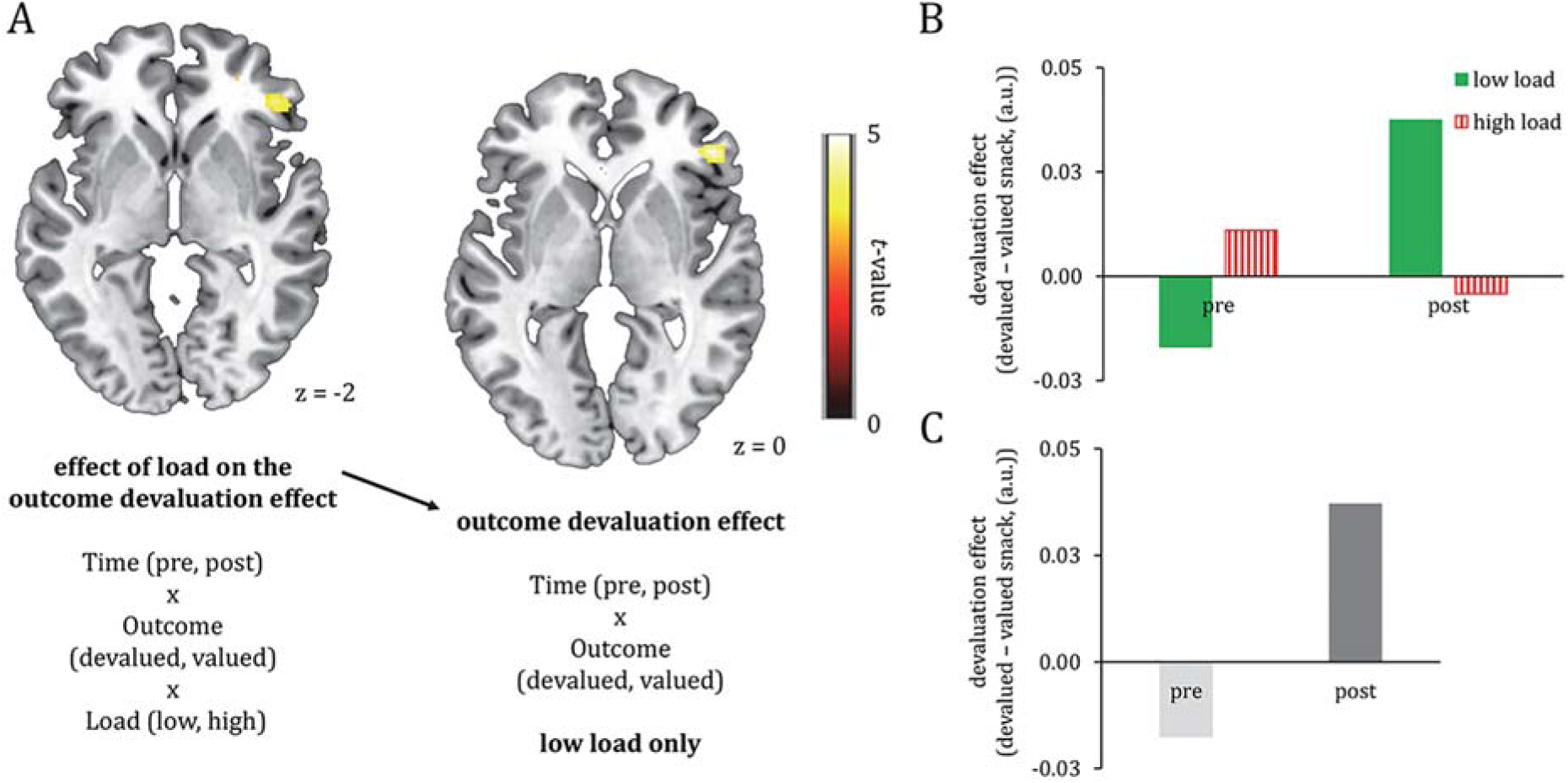
Effect of load on the outcome devaluation effect (Panel A, left: Time (pre,post) x Reward (devalued, valued) x Load (low, high)) and the outcome devaluation effect for the low load conditions only (Panel A, right: two way interaction effect of Time (pre,post) x Reward (devalued, valued)). Both contrasts showed responses of the rIFG. Under high load, the rIFG did not respond, suggesting that distraction tended to disrupt activation of this region. Bar graphs show parameter estimates in arbitrary units (a.u.) for the effect of load on the outcome devaluation effect (Panel B) and the outcome devaluation effect for low load only (Panel C). The bar graphs are shown for illustrative purposes only, therefore error bars and p-values were omitted. All parametric maps are depicted at a *p* < 0.001, uncorrected threshold.

### fMRI results: distraction-related functional connectivity analysis

In a secondary analysis, we performed a gPPI analysis to investigate distraction-related (high-low) functional connectivity during goal-directed effort for food rewards. As a seed we used the region in the rIFG that showed an effect of distraction on goal-directed responses for food reward (Fig. 5, (top left)), extracted at cluster-defining threshold *p* < 0.001 (uncorrected). The rIFG seed region showed increased functional connectivity with bilateral putamen for valued versus devalued food rewards (Time (post>pre) x Reward (valued>devalued) x Load (low>high) interaction contrast). The left region was significant after small volume correction using our anatomically-defined fronto-striatal mask from the Hammersmith atlas (see *Methods*; [-28, −12, 4], *pFWE*(cluster) = 0.004, *t* = 5.16, k = 300, Fig. 5), whereas the right region showed a trend ([22, 6, −8], *pFWE*(cluster) = 0.07, *t* = 4.14, k = 129, Fig. 5). We further investigated this three-way interaction by assessing the Time (post>pre) x Reward (valued>devalued) interaction effect separately for the low and high distraction conditions on the whole brain. Under low, but not high, distraction, the left (but not right) putamen tended to show larger responses for the valued relative to the devalued reward on a *p* < 0.001, uncorrected threshold ([-22, −12, 2], k = 29, *t* = 4.24, *p*FWE(cluster-level) = 0.61). Thus, distraction attenuated connectivity between the right IFG and left putamen during goal-directed effort for food reward.

**Figure 5.**
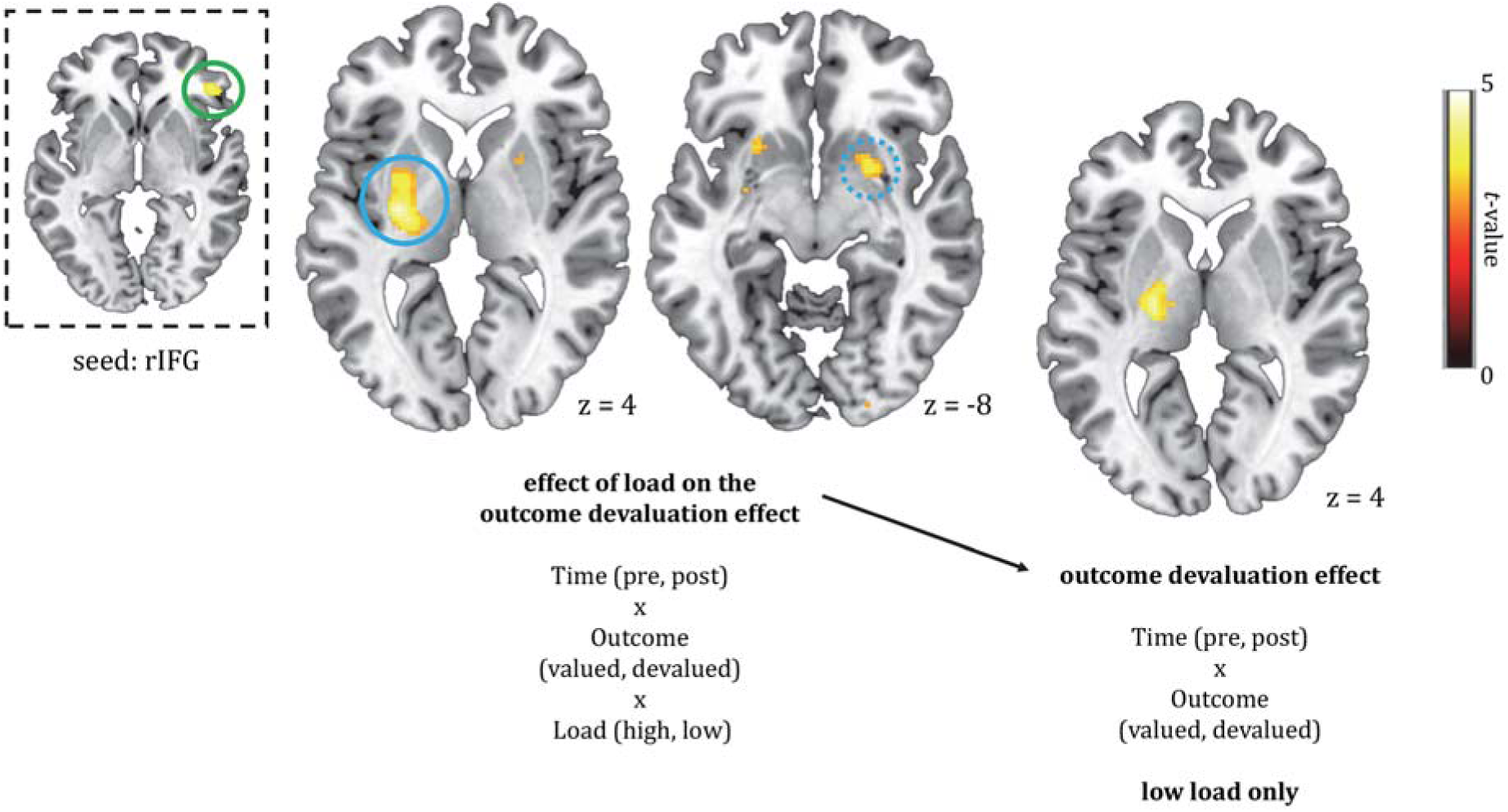
Results of the gPPI analysis with the rIFG seed region in the top left (encircled green, extracted from the Time (post>pre) x Reward (valued>devalued) x Load (low>high) interaction contrast). Shown (encircled blue) is the left [-28,12,4] putamen exhibiting significantly (*p* = 0.004, FWE-corrected at the cluster-level) higher functional connectivity with the seed region under low, relative to high, distraction for valued versus devalued food rewards (effect of load on the outcome devaluation effect). For right [22,6,-8] putamen (encircled blue, dashed), this effect was marginally significant (*p* = 0.07, FWE(cluster)). Further analysis showed this effect tended to be driven by responses of left putamen in the effect of outcome devaluation under low, but not high, distraction ([-22,-12,2], *t* = 4.24, k = 29, *p* < 0.001, uncorrected). All parametric maps are depicted at a *p* < 0.001, uncorrected threshold. For whole-brain (FWE < 0.05) corrected effects see Table 4.

### Brain-behavior correlations

Despite the absence of behavioral effects of distraction on button presses (i.e. effort) for devalued versus valued reward outcomes across the group, exploratory between-subject analyses showed large individual variability in effort that was meaningfully related to brain responses. Specifically, the interaction effect (Load x Reward x Time) in the rIFG co-varied with the same interaction effect on the behavioral effort measure. That is, distraction-related rIFG decreases were associated with continued button presses for food reward after devaluation (four-way interaction effect: *F*(1,36) = 6.69, *p* = 0.01, *r* = −0.40). Further analysis showed this relation was present after the outcome devaluation phase (three-way interaction effect, post: *F*(1,36) = 5.54, *p* = 0.02, *r* = −0.37), but not prior to the devaluation (three-way interaction effect, pre: *F*(1,39) = 2.83, *p* = 0.10, *r* = −0.27). Thus, distraction-induced reductions in rIFG processing were associated with increased effort for food reward after outcome devaluation (Fig. 6).

**Figure 6.**
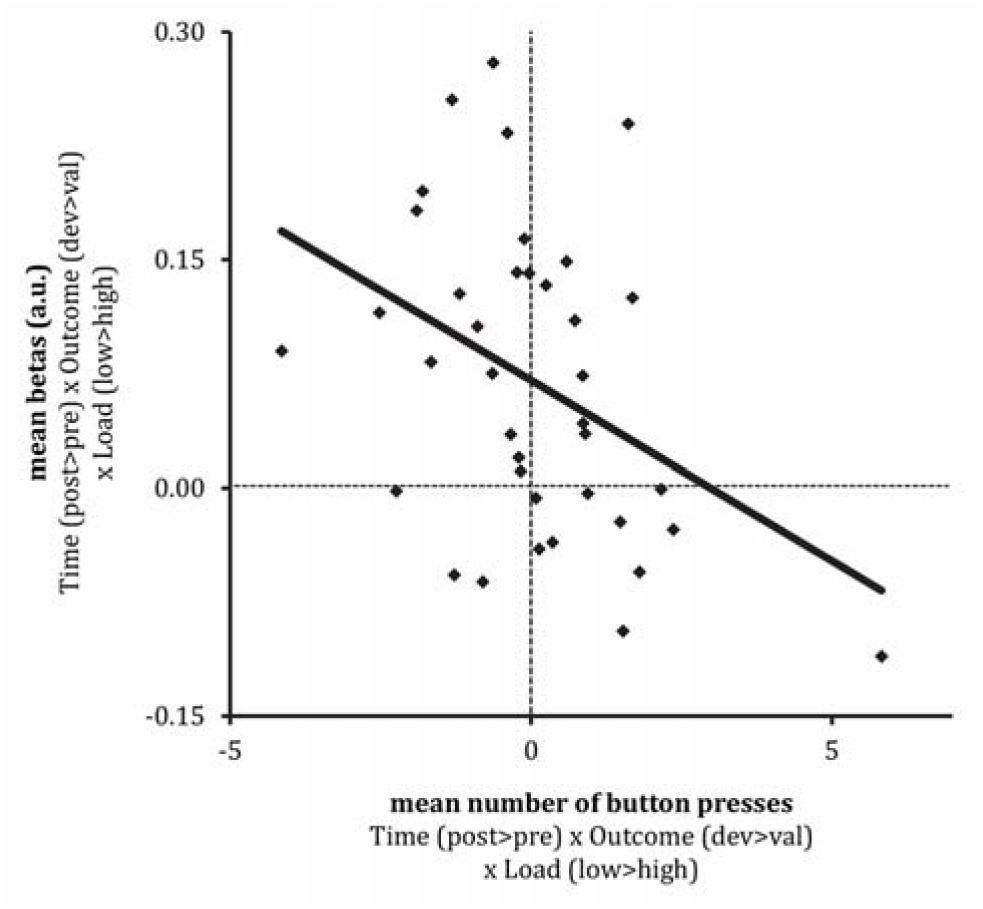
Brain-behavior correlation for the relation between the distraction-sensitive rIFG responses for devalued versus valued food rewards (post-devaluation (devalued reward (low-high distraction) – valued reward (low-high distraction)) – pre-devaluation (devalued reward (low-high distraction) – valued reward (low-high distraction)), and the same effect for continued button presses (effort). The correlation is significant at the highest level (Time (post-pre devaluation) x Reward (devalued-valued reward) x Load (low-high distraction), *r* = 0.040, *p* = 0.014). Less distraction-induced goal-directed rIFG responses relate to continued button presses for food reward. Mean rIFG betas are presented in arbitrary units (a.u.).

To investigate whether the effect of distraction on connectivity between the rIFG and putamen correlated with effort, we used a similar approach. This correlation was not significant.

### Additional results: liking ratings

Similar to the results of the wanting ratings, the mean liking ratings changed significantly over time for the devalued, but not for the valued, reward (Reward (valued, devalued) x Time (t_0_, t_1_, t_2_, t_3_): *F*(1,35) = 18.13, *p* < 0.001, Table 3). Main effects of Time (*F*(1,35) = 24.54, *p* < 0.001) and Reward (*F*(1,37) = 12.83, *p* = 0.001) were significant, reflecting decreased liking for both rewards after the devaluation, and the devalued reward was generally less liked than the valued reward. At baseline (t_0_), liking did not to differ significantly between the rewards (*t*(1,37) = −2.00, *p* = 0.05).

### Additional results: hunger, fullness and thirst ratings

Analysis of the self-reported hunger, fullness, and thirst ratings showed main effects of Time (see Table 3 for means per time point and *F* and *p* statistics). A paired samples t-test using the mean ratings pre-(t_0_, t_1_, t_2_, t_3_) versus post-devaluation (t_4_, t_5_, t_6_, t_7_), showed participants’ hunger decreased, and their fullness increased significantly after the devaluation (mean (± SEM) hunger pre: 7.0(0.3), mean (± SEM) hunger post: 3.5(0.4), *t*(1,37) = 10.82, *p* < 0.001; mean (± SEM) fullness pre: 1.8(0.2), mean (± SEM) fullness post: 6.1(0.3), *t*(1,37) = −14.91, *p* < 0.001). As anticipated, thirst ratings did not change significantly after the devaluation (*t*(1,37) = .798, *p* < 0.43. Thus, participants were successfully satiated on one of the snacks after the outcome devaluation phase.

### Additional results: hierarchical regression

To determine effects of individual differences, we performed two hierarchical regression (using backward elimination) analyses. We used the effect of distraction on processing of the rIFG, and on connectivity between rIFG and left putamen during goal-directed control as dependent variables (interaction effect of Load x Reward x Time for the rIFG, and the psychophysiological interaction effect of Load x Reward x Time for left putamen with the rIFG seed). The independent variables were: wanting (first level), liking, performance, hunger, fullness, and thirst ratings, and the questionnaires (second level) as described in the Methods section. The significance level was Bonferroni-corrected for the two outcome measures and set to α = 0.025. Both regression analyses did not result in significant models (all *p* > 0.025). Thus, there were no individual factors predicting the effect of distraction on goal-directed control in the rIFG or distraction-related connectivity between the rIFG (seed) and left putamen.

## Discussion

To study the effect of distraction on effort for valued and devalued food reward, we used a food-related outcome devaluation paradigm, and manipulated distraction using a categorical visual detection task of varying attentional load. We hypothesized that distraction would disrupt goal-directed control for food reward, by acting on the vmPFC or other fronto-striatal regions (see <u>preregistration</u>).

As a result of outcome devaluation, we expected to find more activation in the vmPFC for valued than devalued food reward, in line with its role in value-based decision making (Valentin et al., 2007; Tanaka et al., 2008; Gläscher et al., 2009; Rangel, 2013). However, the vmPFC did not show this effect and, if anything, the reverse effect (devalued > valued at *p* < 0.001 uncorrected) occurred. Instead, we found significantly more activation in the ACC after outcome devaluation (valued > devalued). This is likely due to the nature of the task, which was rather different from the instrumental choice paradigms used by aforementioned researchers. For the current research question, both experimental design and ecological-validity required the distraction and food-choice manipulations to co-occur for a continuous time. Repeated button presses, i.e. exerting effort, to obtain the food rewards during high or low distraction therefore offered a better manipulation than a single, binary choice paradigm. ACC responses have been associated with effort-based cost-benefit analyses and value-based effort allocation (Kurniawan et al., 2013; Shenhav et al., 2013; Vassena et al., 2017), i.e. computations that are likely to be enhanced for valued versus devalued food rewards.

We were mainly interested in the effects of distraction on responses to valued versus devalued food rewards. However, neither the ACC nor the vmPFC showed this interaction effect. Instead, high – versus low – distraction reduced goal-directed responses for devalued versus valued food rewards in the rIFG. Based on previous studies, we did not specifically anticipate finding this region for the effect of distraction on goal-directed responses. Hence, it did not survive multiple comparison correction in our large search volume of fronto-striatal regions. Interestingly, however, the effect of distraction on rIFG responses correlated significantly with the same effect in our behavioral effort measure (repeated button presses). Under low, but not high, distraction, larger responses of the rIFG related to less repeated button presses for the devalued versus valued food reward. Activation of the right IFG has been strongly associated with response inhibition (Aron et al., 2003; Zandbelt and Vink, 2010). Therefore, this result might reflect worse response inhibition in response to the devalued versus valued food reward under distraction, in line with its negative association with the number of button presses. We speculate that our participants had to suppress their automatic tendency to press for the (now) devalued reward, as their focus was on performing the visual detection task. Our results indicate that for participants with relatively less rIFG activation under high versus low distraction, more effort was exerted for the devalued reward, which suggests worse response inhibition in these individuals.

Furthermore, under high distraction, we found weaker connectivity between the rIFG and left putamen for valued versus devalued food rewards. Like the rIFG, the putamen has been associated with successful response inhibition (Zandbelt and Vink, 2010). This result therefore supports the idea that distraction during repeated button presses for food rewards attenuates the mechanism underlying successful response inhibition.

In contrast to these significant effects of distraction on rIFG-putamen connectivity across the group, our rIFG activity effects were only significant in a between-subject correlation with behavior. This shows that there is substantial individual variation in the effects of distraction on food (reward)-related processing. These individual differences are in line with our previous study, in which distraction during food consumption attenuated taste-related processes in the insula, but only in relation to increases in food intake (<u>Duif et al., submitted</u>). Future studies should investigate the relative contribution of impairments in taste processing versus response inhibition in distracted (over)eating in real life, and the individual susceptibility to distraction during food-related behavior.

Taken together, distraction during effort for food rewards following sensory-specific satiation tends to attenuate responses of the rIFG in individuals that are more susceptible to the distraction manipulation in terms of their behavioral responses, and weakens its functional connectivity with putamen. Both the rIFG and the putamen have been implicated in response inhibition. Moreover, distraction-induced reductions in rIFG signal were indeed related to continued button presses for devalued food reward. Therefore, distraction during food-related effort seems to attenuate response inhibition. Interestingly, impaired response inhibition is also highly implicated in overweight and obesity (Stice and Burger, 2019). Further research should focus on the importance of the (susceptibility for the) effect of attention/distraction on response inhibition in overweight individuals.

## Acknowledgements

We would like to thank Britt Lambregts, Marlou Lasschuijt and Lieke van Lieshout for their help in conducting the study.

